# Alteration in Gut Microbiome and Intestinal Barrier Function caused by Efavirenz versus Dolutegravir Treatments in Mice

**DOI:** 10.1101/2023.11.01.565232

**Authors:** Qin Tang, Hai-en Pan, Zhe Liu, Zhou Deng, Qing-Yang Zhong, Huan-huan Cao, Jun-yan Ye, Jin Li, Xue-ying Li, Hai-peng Zhu, Song Chen

## Abstract

Dolutegravir (DTG) is replacing efavirenz (EFV) as first-line antiretroviral therapy because of its better tolerance. However, DTG cause similar, but milder, gastrointestinal and neurological side effects as EFV does. We speculated that impaired gut barrier function contributes to their side effects. For this purpose, the mice were intragastrically administered EFV, DTG, or vehicle for 60 consecutive days. The plasma levels of FITC-dextran were determined to evaluate gut barrier integrity. Colonic contents were collected for 16S rRNA sequencing. Adipose, liver, ileum, and colon tissues were collected for pathological examination, and intestinal zona occludens-1 (ZO-1) immunofluorescence staining and goblet cell staining were performed. We found that EFV significantly retarded body weight gain, decreased glucose uptake, and caused lipodystrophy and hepatocyte necrosis. EFV also decreased species richness of gut microbiota, increased Verrucomicrobia and Proteobacteria, and decreased Patescibacteria and Cyanobacteria. Moreover, it caused crypt damage, goblet cell loss, reduced ZO-1 expression, impaired gut barrier function, and suppressed expressions of Pdha1 and Ndufv1. Interestingly, DTG impaired barrier function similar to EFV, but the impairment was milder. DTC also inhibited MPC1, MPC2, and Pdha1 expression. Our results suggest a link between abnormal energy metabolism, impaired gut barrier integrity and side effects of EFV and DTG.

## 1. Introduction

Efavirenz (EFV) is a non-nucleoside reverse transcriptase inhibitor, which is recommended as the preferred first-line antiretroviral therapy (ART), in combination with tenofovir disoproxil fumarate and emtricitabine or lamivudine, for HIV-1 treatment in the 2013 World Health Organization (WHO) guidelines. However, EFV can cause a high prevalence of neurocognitive disorders among asymptomatic patients. The neuropsychiatric side effects, including vivid dreams, dizziness, insomnia, decreased concentration, anxiety, and depression, have been described in over 50% of EFV-treated patients ^1–3^, as well as remarkable limb fat loss ^4^. Therefore, in 2018, dolutegravir (DTG), a potent and well-tolerated integrase inhibitor, replaced EFV according to WHO recommendations ^5^. However, because of its high antiviral effect, low pill burden, relatively stable plasma concentration, and even better T-cells recovery^6^, EFV is still administered to millions of patients. Although less common, DTG could also induce similar neuropsychiatric side effects as EFV. Therefore, it is necessary to investigate the pathogenesis of adverse effects of EFV and DTG.

The gut microbiota plays critical role in neuropsychiatric diseases, including autism, mood disorders, depression ^7^, schizophrenia ^8^, Parkinson’s disease ^9^, and Alzheimer’s disease ^10,11^. Therefore, although EFV exerts a direct impact on neurons and glial cells ^12^, we speculate that gut bacterial translocation contributes to the side effects of EFV and DTG. This hypothesis is supported by the upregulation of nuclear factor-kappa B (NF-κB) in EFV-treated individuals ^13^. Therefore, we compared the impact of EFV and DTG on gut microbiota, as well as on the intestinal barrier function. Moreover, there is no evidence of a direct effect of EFV or DTG on the microbiota. Based on the well-recognized mitochondrial toxicity of EFV and DTG ^14–17^ and the knowledge that mitochondrial pyruvate metabolism is necessary and sufficient to maintain the proliferation of intestinal stem cells ^18,19^, we deduced that pyruvate metabolism might contribute to the EFV and DTG caused gut dysbiosis and impairment of gut barrier function.

## 2. Materials and methods

### 2.1. Animals

C57BL/6J mice (7 weeks of age) were obtained from the Guangdong Medical Laboratory Animal Center (GDMLAC) and maintained under specific pathogen-free conditions. All study animals were housed in sterile cages in a room maintained at 23 ± 2℃ with an average humidity of 60 ± 10% and alternating 12-h light and dark cycles. The mice were supplied with water and rodent chow ad libitum. The protocols were approved by the Animal Experimental Ethical Committee of GDMLAC) (No. 2018007). All related procedures were performed in accordance with the recommendations of the NIH Guide for the Care and Use of Laboratory Animals ^20^.

Following a one-week acclimation period, a total of 36 mice were randomly assigned to the DTG, EFV, and normal control (NC) groups, with six males and six females in each group. Animals in the DTG and EFV groups were intragastrically administrated 20.6 mg/kg DTG (equivalent to an adult dose of 100 mg/day) or 123.3 mg/kg EFV (equivalent to 600 mg/day), respectively. DTG and EFV tablets were purchased from Glaxo Operations UK Ltd. or Shanghai Desano Pharmaceuticals Co., Ltd., respectively. Tablets were minced and suspended in 0.5% sodium carboxymethyl cellulose (Tianjin FUCHEN Chemical Reagent Factory, Tianjin, China). Animals in NC group were administered an equal volume (10 mL/kg) of vehicle as control. All treatments were performed once daily for 60 consecutive days.

### 2.2. Fasting glucose level and oral glucose tolerance test (OGTT)

Mice were transferred to cages with fresh bedding and fasted overnight before testing (16 h) before performing the OGTT while ensuring that the mice had access to drinking water. A glucose load of 2 g/kg body weight was administered by intragastric gavage. Blood samples were collected from the tail vein at 0, 15, 30, 60, and 120 min after glucose administration. Blood glucose levels were measured using an OneTouch Ultra system (Johnson & Johnson, New Brunswick, NJ, USA).

### 2.3. Gut leakage measurement

Gut permeability was determined using the fluorescein isothiocyanate (FITC)-dextran assay. FITC-dextran is a nonabsorbable molecule with average molecular weight of 3000-5000 (Sigma-Aldrich, St. Louis, MO, USA). The FITC-dextran was diluted to 100 mg/mL with PBS and mice received a gavage of 0.6 mg/g once after a 4-h fast the day before termination of treatment. After 1 h, blood was collected and anticoagulated, plasma samples were separated, and immediately subjected to FITC-dextran detection. To generate a standard curve, unused FITC-dextran (100 mg/mL) was serially diluted with water. The fluorescence intensity values were read using a Varioskan LUX multimode microplate reader (Thermo Fisher Scientific, Waltham, MA, USA) with wavelengths of 485 (excitation) and 528 (emission). The concentration of FITC-dextran was divided by the weight to normalize the values.

### 2.4. Tissue sample collection and pathological examination

After euthanasia by cervical dislocation, liver, white adipose tissue, jejunum, ileum, and colon were removed, immersed in 10% buffered formalin for 48 h, processed for paraffin embedding, sectioned into 3-μm-thick sections, and stained using hematoxylin and eosin (H&E). Periodic acid-Schiff (PAS)/hematoxylin staining was performed on the goblet cells. Light microscopy images were obtained and analyzed using a DMR microscope system (Leica Microsystems, Wetzlar, Germany). Histological evaluation was performed by a pathologist blinded to the identity of the sample. The number of goblet cells per villus was counted. For each sample, at least ten complete contiguous crypts were analyzed. Images were analyzed using ImageJ software (NIH, Bethesda, MD, USA).

### 2.5. Immunofluorescence analysis

After deparaffinization and antigen retrieval (EDTA 8.0), immunofluorescence detection of the tight junction protein zona occludens-1 (ZO-1) was performed with rabbit anti ZO-1 polyantibody (Proteintech, Wuhan, China) incubation at 4℃ overnight, followed with Alexa Fluor 594 conjugated goat anti-rabbit lgG (H+L) (ZSGB-BIO, Beijing, China) incubation at 37℃ for 1 h. After adding coverslips with anti-fade mountant containing 4’,6-diamidino-2-phenylindole (DAPI; ZSGB-BIO), the slides were observed and photographed using a TCS SP8 laser confocal microscope (Leica Microsystems) with LAS X Life Science Microscope Software.

### 2.6. 16S rRNA sequencing

Colonic contents were collected into sterilized tubes, immediately frozen in liquid nitrogen, and maintained at -80℃. The DNA procedures were performed as previously described ^21^. Briefly, total genomic DNA from the samples was extracted using the CTAB/SDS method. DNA concentration and purity were monitored by1% agarose gel electrophoresis. According to the concentration, DNA was diluted to 1 ng/μL using sterile water. Gene amplification, cloning, and sequencing of the bacterial 16S rRNA PCR products were conducted in a laboratory maintained by Applied Protein Technology Co. Ltd. (Shanghai, China). PCR amplification of the V5-V4 regions of the bacterial 16S rRNA gene ^22,23^ was conducted using universal primers targeting 16S V3-V4 (341F-806R), 18S V9 (1380F-1510R), and ITS1 (ITS1F-ITS2R) which incorporate unique sample barcode sequences.

To generate an amplicon library, 20 ng of each genomic DNA sample, 1.25 U of Taq DNA polymerase, 5 μL of 10× Ex Taq buffer, 10 mM dNTPs (all reagents from TaKaRa Biotechnology Co., Ltd., Dalian, China), and 40 pmol of primer mix were added to a 50-μL reaction mixture. The PCR conditions were as follows: a 5 min initial denaturation at 95℃, 28 cycles of denaturation at 95℃ (30 s), annealing at 55℃ (30 s), and elongation at 72℃ (45 s), followed by a final extension at 72℃ for 7 min. The same volume of 1× loading buffer containing SYB green with PCR products and operate electrophoresis on 2% agarose gel for detection. Samples with a bright main strip between 400-450bp were chosen for further experiments. The PCR products were mixed at equidensity ratios. The PCR products were purified using an AxyPrepDNA Gel Extraction Kit (AXYGEN Biosciences, Union City, CA, USA). Sequencing libraries were generated using the NEBNext Ultra™ DNA Library Prep Kit for Illumina (NEB, Ipswich, MA, USA) following the manufacturer’s recommendations, and index codes were added. Library quality was assessed on a Qubit 2.0 Fluorometer (Thermo Fisher Scientific) and a Bioanalyzer 2100 system (Agilent, San Diego, CA, USA). Finally, the library was sequenced on an Illumina Miseq/HiSeq2500 platform and 250bp/300bp paired-end reads were generated, and the sequencing data were deposited in the NCBI Sequence Read Archive (SRA) database with accession number accession number PRJNA900705.

For bioinformatics analysis, paired-end reads from the original DNA fragments were merged using FLASH, a very fast and accurate analysis tool that was designed to merge paired-end reads when at least some of the reads overlapped the read generated from the opposite end of the same DNA fragment. Paired-end reads were assigned to each sample according to their unique barcodes.

Sequence analysis was performed using the UPARSE software package with UPARSE-OTU and UPARSE-OTUref algorithms. In-house Perl scripts were used to analyze alpha (within samples) and beta (among samples) diversity. Sequences with ≥97% similarity were assigned to the same operational taxonomic unit (OTU). We selected representative sequences for each OTU and used the RDP classifier to annotate the taxonomic information for each representative sequence. To compute alpha diversity, we rarified the OTU table and calculated three metrics. Chao1 estimates the species abundance, Observed Species estimates the number of unique OTUs found in each sample, and the Shannon index. Rarefaction curves were generated based on these metrics. A graphical representation of the relative abundance of bacterial diversity from phylum to species can be visualized using the Krona chart. We used the unweighted UniFrac distance for Principal Coordinate Analysis (PCoA). PCoA helps obtain principal coordinates and visualize them from complex multidimensional data.

To confirm differences in the abundances of individual taxa between the two groups, STAMP software was used. Linear discriminant analysis Effect Size (LEfSe) was used for the quantitative analysis of biomarkers within different groups and was visualized with a cladogram. This method was designed to analyze data in which the number of species was much higher than the number of samples, and to provide biological class explanations to establish statistical significance, biological consistency, and effect-size estimation of predicted biomarkers. A graphical representation of the relative abundance of bacterial diversity from phylum to species was visualized using the Krona chart. We also used PICRUSt to perform functional classification of the Kyoto Encyclopedia of Genes and Genomes (KEGG) Orthology (KOs) and Clusters of Orthologous Groups (COGs) ^24^.

### 2.7. RNA exaction, reverse transcription, and quantitative Real-Time PCR

The aliquots of intestinal samples were also frozen in liquid nitrogen and maintained at -80℃. RNA was extracted with TRIzol reagent (Thermo Fisher Scientific, Shanghai, China), reverse transcribed using the PrimeScript RT Master Mix (TaKaRa Biomedical Technology Co., Ltd.), and quantified by real-time PCR with TB Green Premix Ex Taq (TaKaRa Biomedical Technology Co., Ltd.) and ABI 7500 (Applied Biosystems Inc., Foster City, CA, USA). The relative expression levels of the analyzed genes were calculated using the 2^-ΔΔCt^ method. Primer sequences are provided in **Supplementary Table S1**.

### 2.8. Statistical analysis

The results are presented as mean ± SD values. Multiple group comparisons were assessed through one-way analysis of variance (ANOVA) with the least significant difference (LSD) or Kruskal-Wallis multi-comparisons test using IBM SPSS software (22.0). Statistical significance was set at *P*<0.05.

## 3. Results

### 3.1. EFV causes loss of body weight, reduced glucose uptake, and abnormal histopathology of liver and adipose tissue in mice

Throughout the study, the body weights of all animals increased gradually, except for a transient slight decrease in the EFV-treated mice in the first week. Moreover, the mice that received EFV treatment gained significantly less weight from week 2 and persisted throughout the experiment (**Fig. 1A**). DTG treatment also reduced weight gain slightly compared to the NC group, but not significantly.

**Figure 1.**
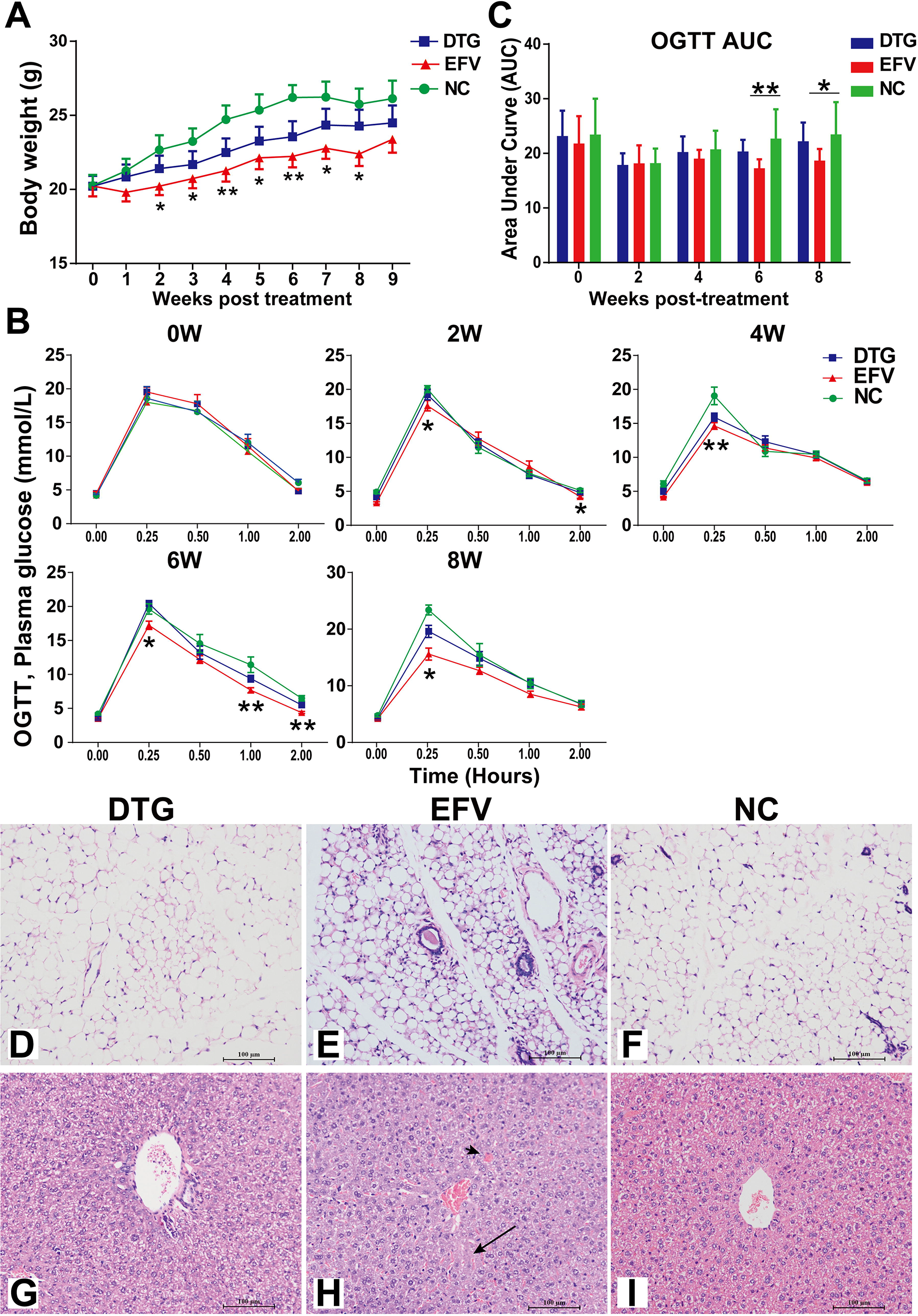
Body weight, oral glucose tolerance test (OGTT), and histopathologic changes in liver and adipose of efavirenz (EFV)-treated and dolutegravir (DTG)-treated mice. (A) Dynamics of body weight, (B) Oral glucose tolerance test (OGTT) results, and (C) the area under curve (AUC) of OGTT; (*Mean ± S.D*). N=12 for each group and \**P*<0.05 or \*\**P*<0.01. (D-F) Representative photomicrographs of adipose tissue from mice in (D) DTG, (E) EFV, and (F) Normal Control (NC) groups, respectively. (G-I) Representative photomicrographs of liver sections from mice in (G) DTG, (H) EFV, and (I) NC groups, respectively. Apoptotic (acidophil) body (short arrow), lobular disarray, and hepatocyte necrosis (long arrow) could be seen in the liver sections of EFV-treated mice. (D-I) Sections were stained with hematoxylin and eosin (H&E).

We also performed a standard OGTT for each animal every 2 weeks. As shown in **Fig. 1B**, fasting glucose levels were not affected by either EFV or DTG treatment throughout the study. In contrast, EFV caused a significant decrease in the peak glucose level 15 min after intragastric glucose injection compared to the NC group. This alteration emerged early (2 weeks) and was maintained throughout the study. EFV may have reduced glucose absorption gradually, which explained the significantly lower area under the curve (AUC) of the OGTT in the EFV group than in the NC group at weeks 6 and 8 (**Fig. 1C**). DTG had less of an impact on glucose levels in the OGTT. Histological examination revealed lipoatrophy in white adipose tissue (**Fig. 1D-F**) and hepatolysis in liver (**Fig. 1G-I**) in EFV-treated mice, recapitulating the commonly reported adverse effects of EFV.

### 3.2. Comparison of EFV and DTG on microbiota community

A total of 3,073,233 high-quality reads with an average length of 422 bp were obtained from the 36 colonic content samples. These reads were combined into 1,375,668 tags, with 38,213 tags per sample. We plotted Rarefaction and Shannon curves to estimate species richness as indices of sequencing depth at the taxonomic level. As shown in **Supplementary Fig. 1A and B,** both the rarefaction and Shannon curves tended to plateau, indicating that the sequencing depth was reasonable and sufficient. A rank-abundance distribution curve for the microbial communities was constructed to visualize species richness and evenness, in which the EFV apparently reduced species richness **(Supplementary Fig. 1C)**. Species accumulation curves (Specaccum) were also plotted to compare the diversity properties of community datasets **(Supplementary Fig. 1D)**.

To assess overall differences in microbial community structure among the groups, we measured ecological parameters based on alpha-diversity (observed species, ACE, and Chao1, Shannon, Simpson, and Phylogenetic diversity (PD) whole tree indices). As shown in **Fig. 2A-2F**, the mean alpha diversity values significantly decreased in the DTG group compared to the NC group (*P*<0.001 for all indices, except Shannon index, for which *P*<0.05). The decrease in the EFV group was even more profound for all indices. In contrast, there was a significant decrease in the Simpson index in the DTG group (*P*<0.05), but not in the EFV group, compared to the NC group.

**Figure 2.**
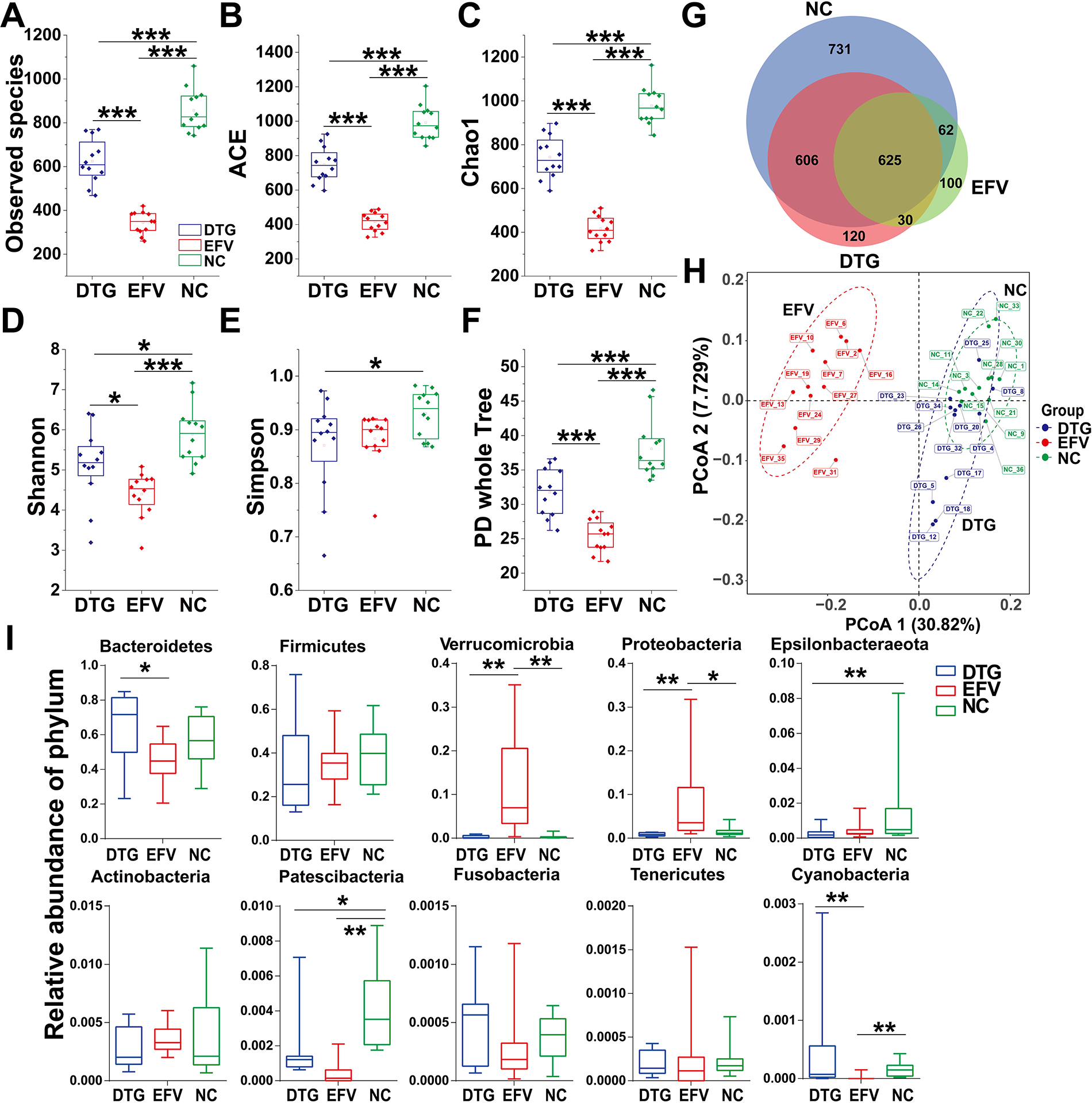
Effects of dolutegravir (DTG) and efavirenz (EFV) on the gut microbial Community distribution. Alpha diversity indices were used to analyse the complexity of gut microbial species of the EFV, DTG, and NC groups. (A) Indices of Observed species, (B) ACE, (C) Chao1, (D) Shannon, (E) Simpson, and (F) PD whole tree are presented with box plots. Moreover, (G) Venn diagram and (H) Unweighted UniFrac-based PCoA plot of the mouse colonic bacterial community showed the diversity among groups. Numbers in brackets represent the contributions of principal components to differences among samples. (I) Comparison of bacterial abundance among DTG, EFV, and NC groups at phylum-level. *N* = 12 for each group. *, ** indicate statistical significance with *P*<0.05 or *P*<0.01 respectively.

We also plotted a Venn diagram showing the number of unique and shared (overlapping regions) OTUs identified by 16S sequencing. There were 731, 120, and 100 unique OTUs in the NC, DTG, and EFV groups, respectively, implying a marked loss of species caused by either DTG or EFV (**Fig. 2G**). To display differences in OTU composition among different samples, PCoA with Weighted UniFrac analysis was used to construct a two-dimensional graph summarizing the factors primarily responsible for this difference. The first coordinate (PcoA 1) elucidated 30.82% of the inter-sample variance. PcoA 2 elucidated 7.729% of differences between the three groups (**Fig. 2H**). The PcoA plot revealed a markedly clustered separation of different groups, and the EFV group separated clearly from the partially overlapped NC and DTG groups, indicating a more severe impact of EFV than that of DTG.

According to the OTU phylum-level annotations, we compared the relative abundances of ten enriched phyla (Bacteroidetes, Firmicutes, Verrucomicrobia, Proteobacteria, Epsilonbacteraeota, Actinobacteria, Patescibacteria, Fusobacteria, Tenericutes, and Cyanobacteria) using the Kruskal-Wallis test. Compared to the NC group, Epsilonbacteraeota and Patescibacteria decreased significantly in the DTG group (*P*<0.01 and *P*<0.05, respectively), whereas Verrucomicrobia and Proteobacteria increased significantly (*P*<0.01 and *P*<0.05, respectively), accompanied by a significant decrease in Patescibacteria and Cyanobacteria abundances (both *P*<0.01) in the EFV group (**Fig. 2I**).

### 3.3. DTG and ETV cause gut dysbiosis

At the genus level, a total of 35 bacteria were identified with significantly different abundances (*P*<0.05) (STAMP, t-test) with a threshold *P*<0.05 in DTG-treated mice compared to NC-treated mice. Ranked by the difference in mean proportions, the five most enriched genera were decreased *other* (belonging to the Lachnospiraceae family, as shown in **Supplementary Fig. 3** in which indicated by letter “*t*”), *uncultured Bacteroidales bacterium, Lachnospiraceae NK4A136 group,* and increased *Alloprevotella, Alistipes* (**Fig. 3A**). Based on PICRUSt, changes in the functional capacity of the gut microbiota were predicted, as indicated by KEGG pathways. A total of four differential KEGG pathways, including increased Digestive System and decreased Cell Motility, Cell Communication, and Sensory System, were enriched (**Fig. 3B**).

**Figure 3.**
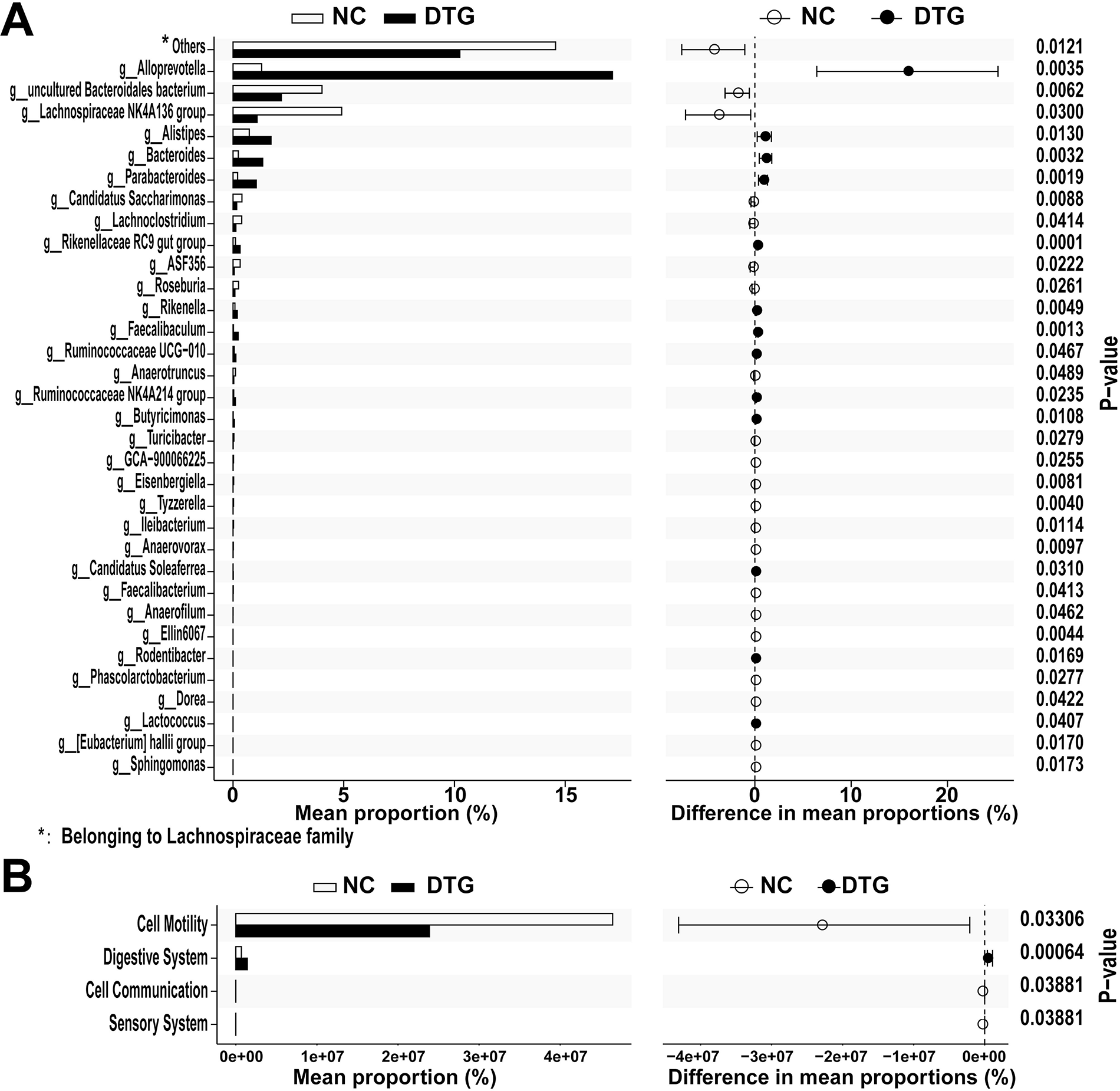
Genera with differential abundance and predicted Kyoto Encyclopedia of Genes and Genomes (KEGG) pathways enriched in DTG group compared to NC group. (A) Relative proportion of genera with differential abundance in the DTG group compared to the NC group. (B) Predicted KEGG pathways with differential relative proportion in comparison between the DTG and NC groups.

EFV caused even more profound alteration in the gut microbiota. At the genus level, a total of 75 bacteria were identified with significantly different abundances (*P*<0.05) (STAMP, t-test) with a threshold *P*<0.05 in EFV-treated mice compared to NC-treated mice. Ranked by the difference in mean proportions, the five most enriched genera were decreased *in uncultured bacteria* (belonging to the Muribaculaceae family)*, Lactobacillus,* and increased *in Bacteroides, Akkermansia,* and *Dubosiella* **(Fig. 4A, Supplementary Fig.4)**. There were 38 predicted KEGG pathways, among which Membrane Transport, Carbohydrate Metabolism, Amino Acid Metabolism, Replication and Repair, and Energy Metabolism were the top five enriched pathways (**Fig. 4B**). As expected, the Nervous System and Neurodegenerative Diseases KEGG pathways were also enriched but ranked only 29^th^ and 30^th^, respectively.

**Figure 4.**
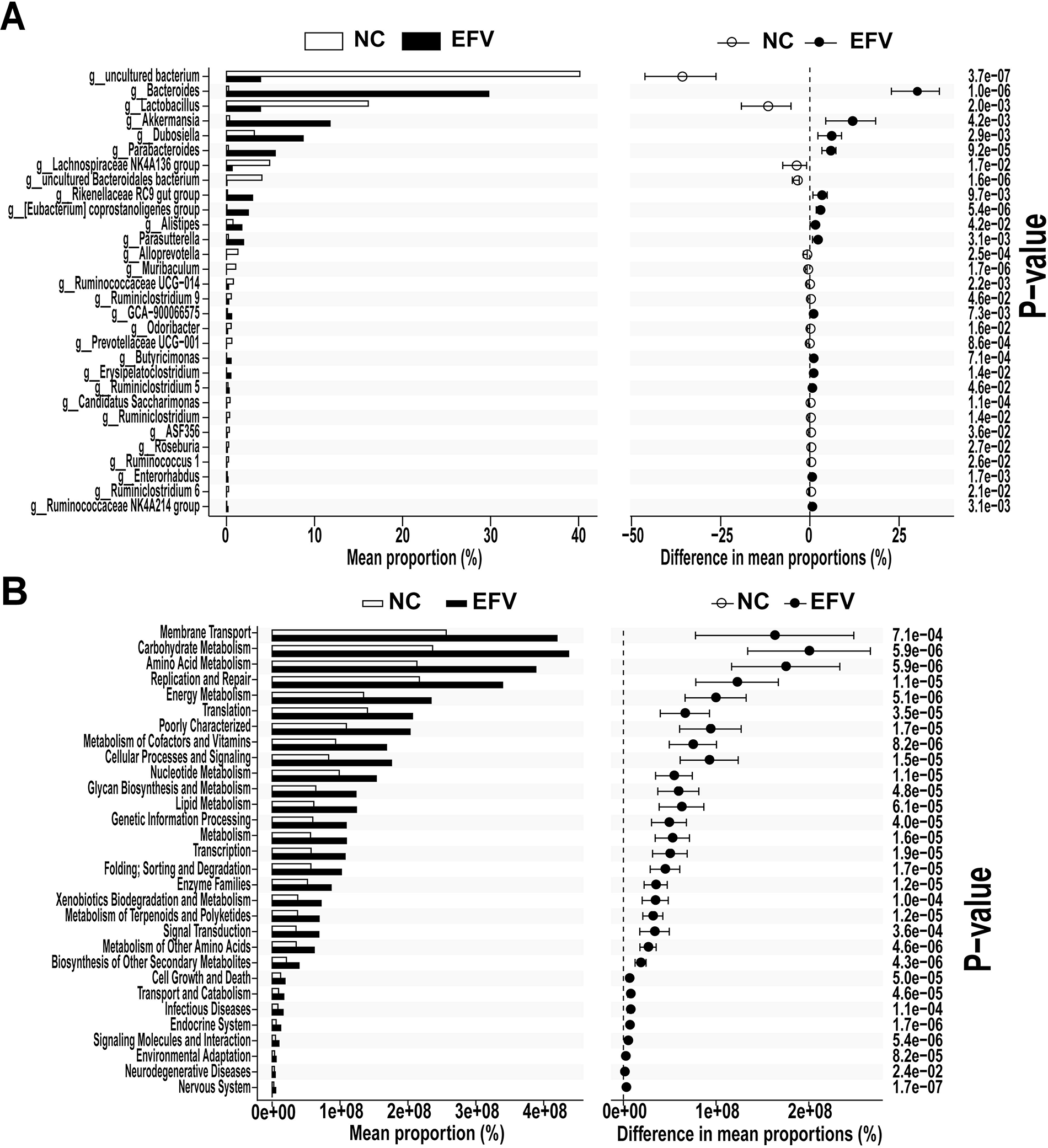
Genera with differential abundance and predicted Kyoto Encyclopedia of Genes and Genomes (KEGG) pathways enriched in EFV group compared to NC group. (A) Relative proportion of genera with differential abundance in the EFV group compared to the NC group. (B) Predicted KEGG pathways with differential relative proportion in comparison between the EFV and NC groups.

We also compared the effects of EFV and DTG on the gut microbiota. As shown in **Supplementary Fig. 2A,** the most remarkable alteration in genus were Muribaculaceae family (including uncultured bacterium, uncultured Bacteroidales bacterium, and muribaculum, indicated by v, w, and x, respectively in **Supplementary Fig. 5**) and KEGG pathways in this comparison were similar to those identified in EFV samples.

### 3.4. Gut barrier function and ileal pathological examination

We further compared the impact of EFV and DTG on gut barrier defects in the FITC-dextran (4.4 kDa) assay. As shown in **Fig. 5A**, both EFV and DTG induced a significant increase in plasma FITC-dextran concentration (both *P*<0.01), indicating increased permeability of the gut barrier. Moreover, the FITC-dextran level was higher in the EFV group than in the DTG group, although the difference was not statistically significant. Using immunofluorescence staining for the intestinal tight-junction molecule ZO-1, we further observed weakened crypt staining in the EFV group (indicated by the long arrow in **Fig. 5B-5D**). Correspondingly, EFV treatment extensively destroyed the crypt structure and induced crypt atrophy and even crypt loss in H&E stained sections (indicated by the short arrow in **Fig. 5E-5G**). EFV treatment also caused a significant loss of goblet cells compared to DTG treatment and a marginally significant loss compared to the NC group with representative images in **Fig. 5H-5J**. The goblet cell counts are shown in columns in **Fig. 5K**.

**Figure 5.**
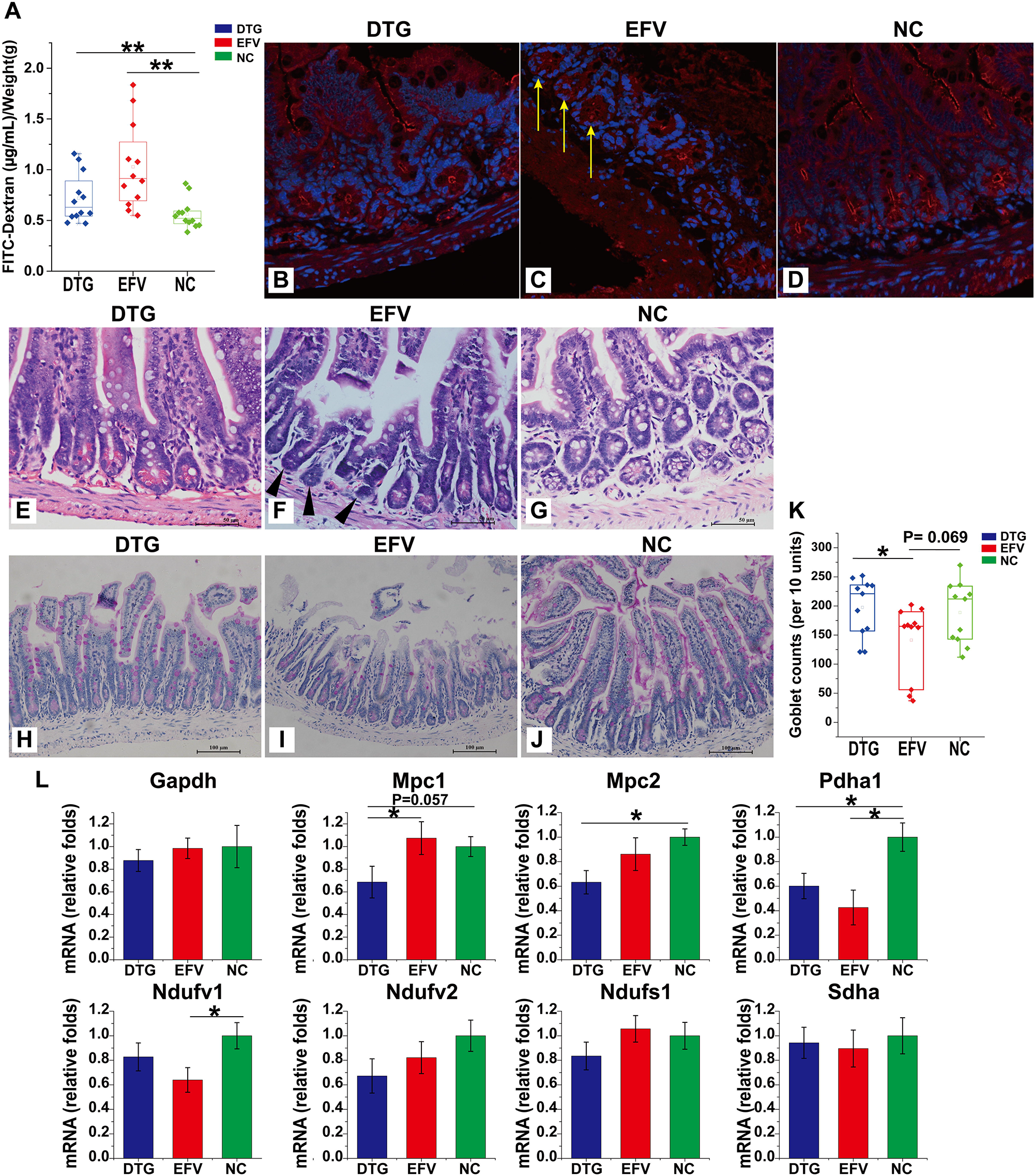
Impaired intestinal barrier function and suppressed expression of pyruvate metabolism-associated genes in the ileum caused by EFV versus DTG. (A) The plasma levels of FITC-dextran indicating gut permeability 1 hour after oral FITC-dextran administration. Values were normalized to body weight. Data were presented as mean ± SD values. *N* = 12 for each group. ** indicate statistical significance with *P*<0.01. (B-D) The representative pictures of immune-fluorescent staining of zona occludens-1 (ZO-1). Ileal samples from mice in (B) DTG, (C) EFV, and (D) NC groups were used. Alexa Fluor 594 (red) and DAPI (blue) was used for ZO-1 and nuclear staining, respectively. (E-G) The representative pictures of ileal histopathologic changes of mice in (E) DTG, (F) EFV, and (G) NC groups with H&E staining. (H-J) The representative pictures of ileal Periodic-Acid-Schiff (PAS) staining indicating goblet cells. (K) Goblet cell counts were shown in columns. Data were presented as mean ± SD values. *N* = 12 for each group.* indicates statistical significance with *P* < 0.05. (L) The relative folds of expression of intestinal pyruvic acid metabolism-associated genes. The mRNA levels were quantified with quantitative PCR analysis and normalized to beta-actin. N = 12 for each group. * indicate statistical significance with *P* < 0.05 compared to NC group.

### 3.5. EFV and DTG alter the expression profile of pyruvate metabolism associated genes

The expression profiles of genes encoding pyruvate metabolism enzymes (mitochondrial pyruvate carrier Mpc1, Mpc2, pyruvate dehydrogenase Pdha1), subunits of mitochondrial complex I (NADH:ubiquinone oxidoreductase) (Ndufv1, Ndufv2, and Ndufs1), complex II (Sdha) of the mitochondrial respiratory chain, and glyceraldehyde 3-phosphate dehydrogenase (Gapdh), which catalyzes the sixth step of glycolysis, were quantified. As shown in **Fig. 5L**, DTG significantly inhibited the expression of Mpc1, Mpc2, and Pdha1, and EFV significantly inhibited Pdha1 and Ndufv1expreesion, while Ndufs1 and Sdha were not significantly affected.

## 4. Discussion

The long-term use of ART drugs has caused an increasing incidence of gastrointestinal, neuropsychiatric, hepatic or metabolic side effects. In this study, we recapitulated weight loss, impaired glucose intake, and histopathologic changes in white adipose and liver tissues in EFV-treated mice. This provided a model to investigate ART-associated adverse effects. As expected, we observed severe impairment of gut barrier function in EFV-treated mice. Considering the well-documented connection between intestinal barrier integrity, bacterial translocation, and neuropsychiatric side effects ^25^, this result provides a novel perspective for understanding the pathogenesis of the adverse effects of EFV and suggests a possible therapy for its treatment. Moreover, unexpectedly, gut barrier integrity was also compromised in DTG treated mice, although it was milder than that in EFV-treated mice. This suggests a possible explanation for the neuropsychiatric side effects of DTG.

The pathogenesis of gut barrier impairment remains unclear. As the epithelium of the murine small intestine is renewed every 5 days, derived from Lgr5+ stem cells located at the crypt bottoms ^26^, it is reasonable to suggest that the destroyed crypt structure might contribute to impaired gut barrier function. Mitochondrial complex I and pyruvate metabolism maintain the proliferation of intestinal stem cells ^18,19^. Pyruvate is transferred into the inner mitochondrial matrix by Mpc1 and Mpc2 ^27^, converted into acetyl-CoA by the pyruvate dehydrogenase complex, and then subjected to the tricarboxylic acid cycle for oxidative phosphorylation. EFV specifically inhibits complex I by EFV ^28^, whereas DTG inhibits complex IV ^17,26,29^. Consistently, we found that EFV significantly inhibited the expression of Pdha1 and Ndufv1 (encoding the 51 kDa subunit of complex I), while DTG downregulated Mpc1, Mpc2, and Pdha1 expression. These results imply that the abnormal pyruvate metabolism caused by either EFV or DTG might regulate intestinal stem cells and thereby impair barrier function.

The richness of the human gut microbiome is correlated with metabolic markers ^30^. In this study, we observed a significant loss in gut microbiome richness in both the EFV and DTG groups. Consistent with the worse gut leakage, EFV caused more severe richness loss and gut dysbiosis than DTG. However, whether gut dysbiosis contributes to the pathogenesis of EFV-and DTG-associated side effects or is solely the consequence of abnormal mucosal pyruvate metabolism and immunity remains uncertain.

Our research is helpful for understanding the pathogenesis of ETF-and DTG-associated side effects, which could be attributed to pyruvate metabolism, abnormal intestinal stem cell and crypt destruction, compromised barrier function, and bacterial translocation, which has long been associated with neuropsychiatric adverse reactions. Nevertheless, further research is needed to investigate whether gut bacteria penetrate the barrier and destroy the crypt or whether it is shaped by the altered intestinal mucosal environment.

## Abbreviations

ART: Antiretroviral therapy
DTG: Dolutegravir
EFV: Efavirenz
FITC: fluorescein isothiocyanate
Mpc1: mitochondrial pyruvate carrier 1
Mpc2: mitochondrial pyruvate carrier 2
Ndufs1: NADH:ubiquinone oxidoreductase Core Subunit S1
Ndufv1: NADH:ubiquinone oxidoreductase Core Subunit V1
Ndufv2: NADH:ubiquinone oxidoreductase Core Subunit V2
NF-κB: Nuclear factor-kappa B
Pdha1: Pyruvate dehydrogenase
Sdha: Succinate dehydrogenase complex, subunit A
Gapdh: glyceraldehyde 3-phosphate dehydrogenase
ZO-1: Zona occludens-1

## Acknowledgements

We thank the Lingnan Medical Research Center of Guangzhou University of Chinese Medicine for providing some of the laboratory facilities used in this study.

## Financial statement

This work was supported by grants from “Social Science and Technology Development Program of Dongguan” (No. 2020507150119175) and “Social Science and Technology Development Key Program of Dongguan” (No. 20221800905402). The funders had no role in study design, data collection and analysis, decision to publish, or preparation of the manuscript.

## Data availability statement

The sequencing data were deposited in the NCBI Sequence Read Archive (SRA) database with accession number accession number PRJNA900705. The data that support the findings of this study are available from the corresponding author upon reasonable request.

## Subject section and Specified Fields

Bacteriology: Pathogenesis; Immunology: Allergy and inflammation

