## Supplementary Materials for "Alteration in Gut Microbiome and Intestinal Barrier Function caused by Efavirenz versus Dolutegravir Treatments in Mice"

The supplementary file includes:

**Supplementary Table 1.**

**Supplementary Figure 1 to 5.**

**Supplementary Table 1. Sequences of primers.**

|  | **Forward (5’-3’)** | **Reverse(5’-3’)** |
| --- | --- | --- |
| β-actin | GTCCCTCACCCTCCCAAAAG | GCTGCCTCAACACCTCAACCC |
| Gapdh | CATGGCCTTCCGTGTTCCTA | GCGGCACGTCAGATCCA |
| Mpc1 | CGCCCTCTGTTGCTATTCTCTGAC | GGCCGCTTACTCATCTCGTAGTTG |
| Mpc2 | GAAATTGAGGCCGCTTTACAACCAC | GCTAGTCCAGCACACACCAATCC |
| Pdha1 | TTATACGGCGATGGTGCTGCTAATC | GCTGGCTGCTGCTCTCTCAAC |
| Ndufv1 | GTGAGGGCGTTGACTGGATGAAC | GCCTTCTATCTGCTTGCTGATCTCC |
| Ndufv2 | GCTGCTGTGCTTCCAGTCCTG | ACTTCAGCCACCTTGTTCATAGCG |
| Ndufs1 | GTTCTTGCTGACCCACTCGTTCC | ATTGTCTGTGAGGCTCTGCTAATGG |
| Sdha | TGGACATCAAGACTGGCAAGGTTAC | GTAGGAGCGGATAGCAGGAGGTAC |

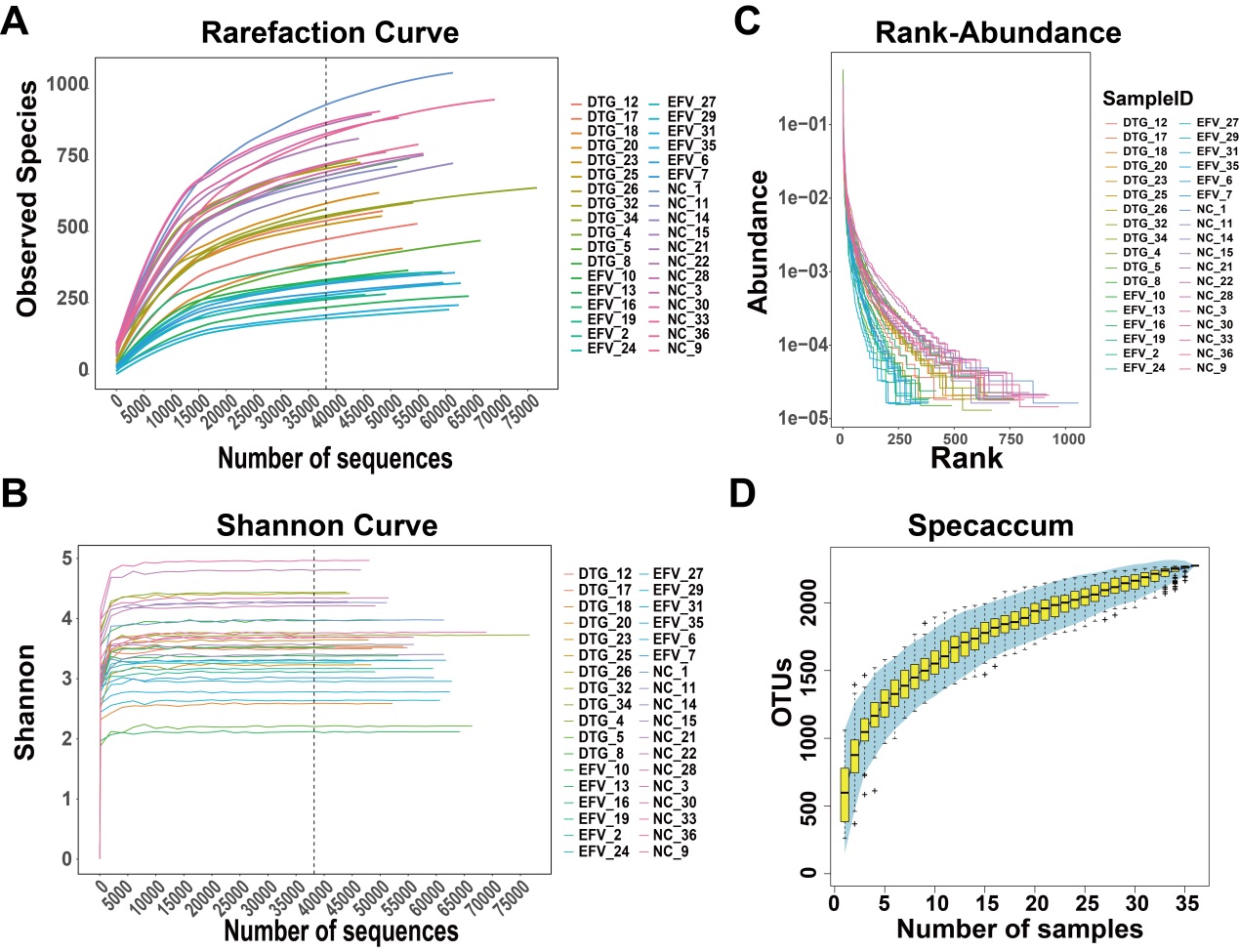

**Supplementary Figure 1. alpha diversity of gut microbiome.**

(A) Refraction curve, (B) Shannon Curve, (C) Rank-Abundance, (D) Specaccum of gut bacterial community of the mouse colonic contents.

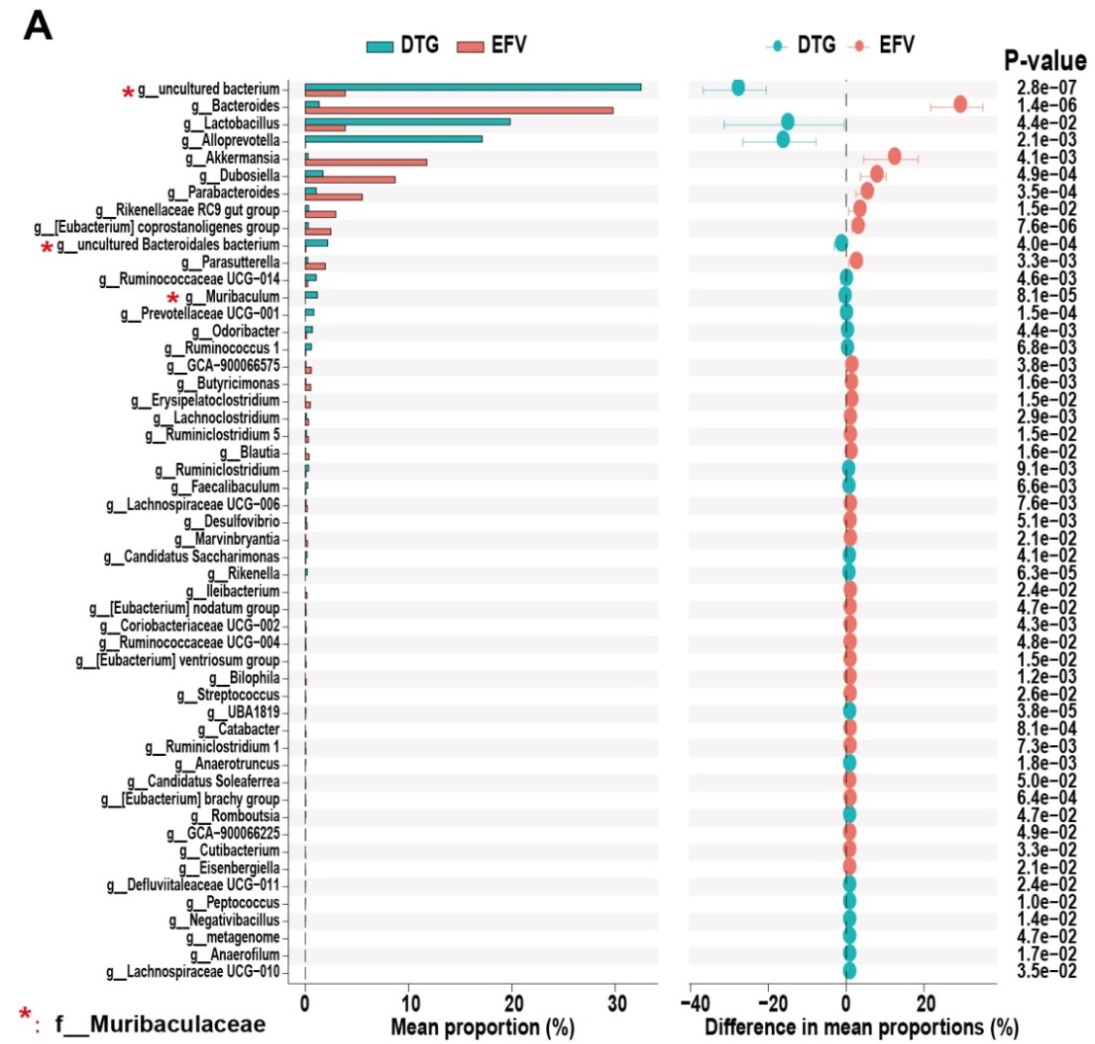

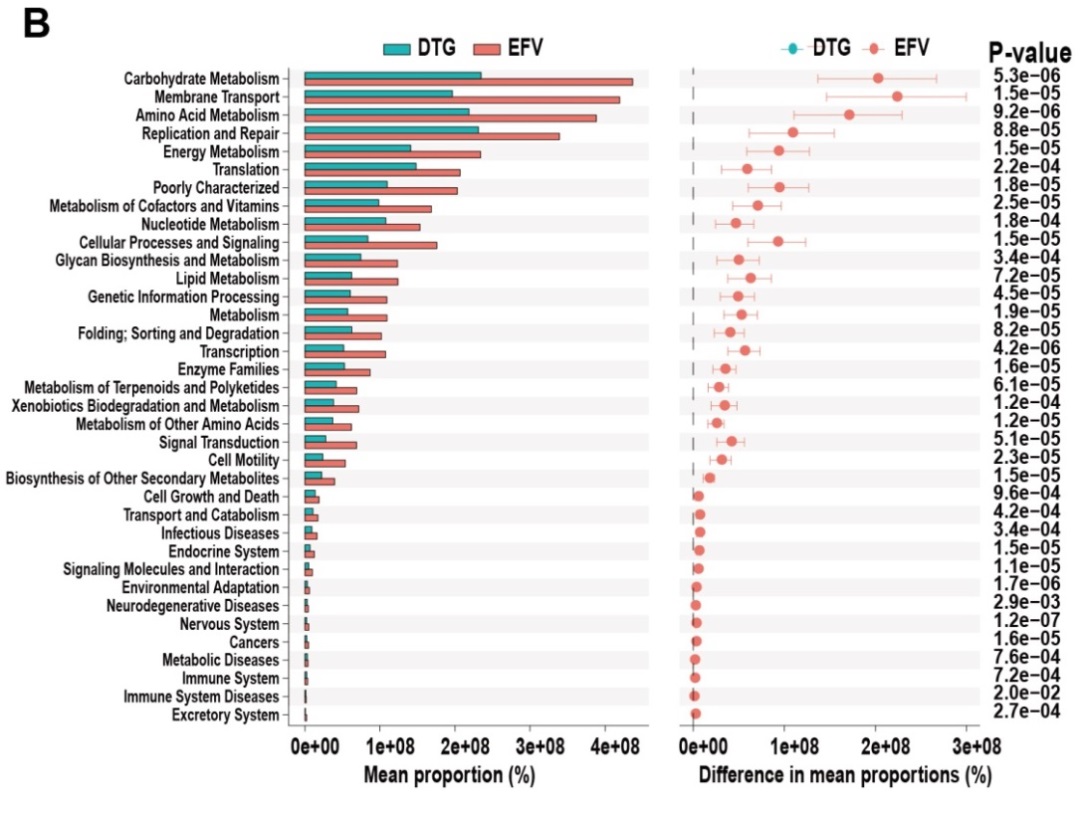

**Supplementary Figure 2. Functional analysis of the gut microbiota using PICRUSt based on the 16S rRNA data**

(A) The abundance of the different phyla Columns show 3 differential abundant categories of KEGG pathways. (*：*P*< 0.05) (B) The subdivision of KEGG pathways with differential relative abundance are shown in a bar plot. (*：*P*< 0.05)

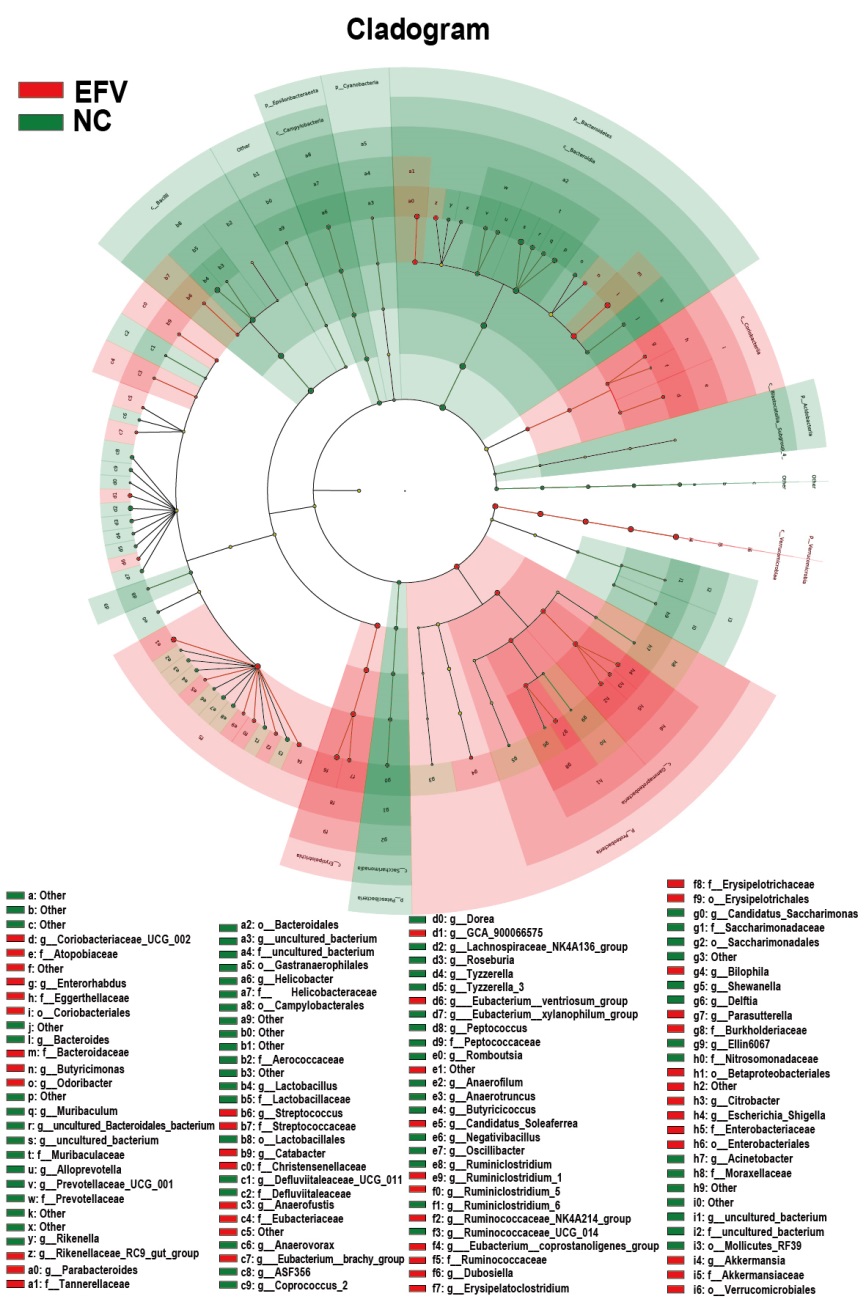

**Supplementary Figure 3. Cladogram of linear discriminant analysis effect size (LEfSe) showing the microbiota with different abundances between EFV and NC groups.**

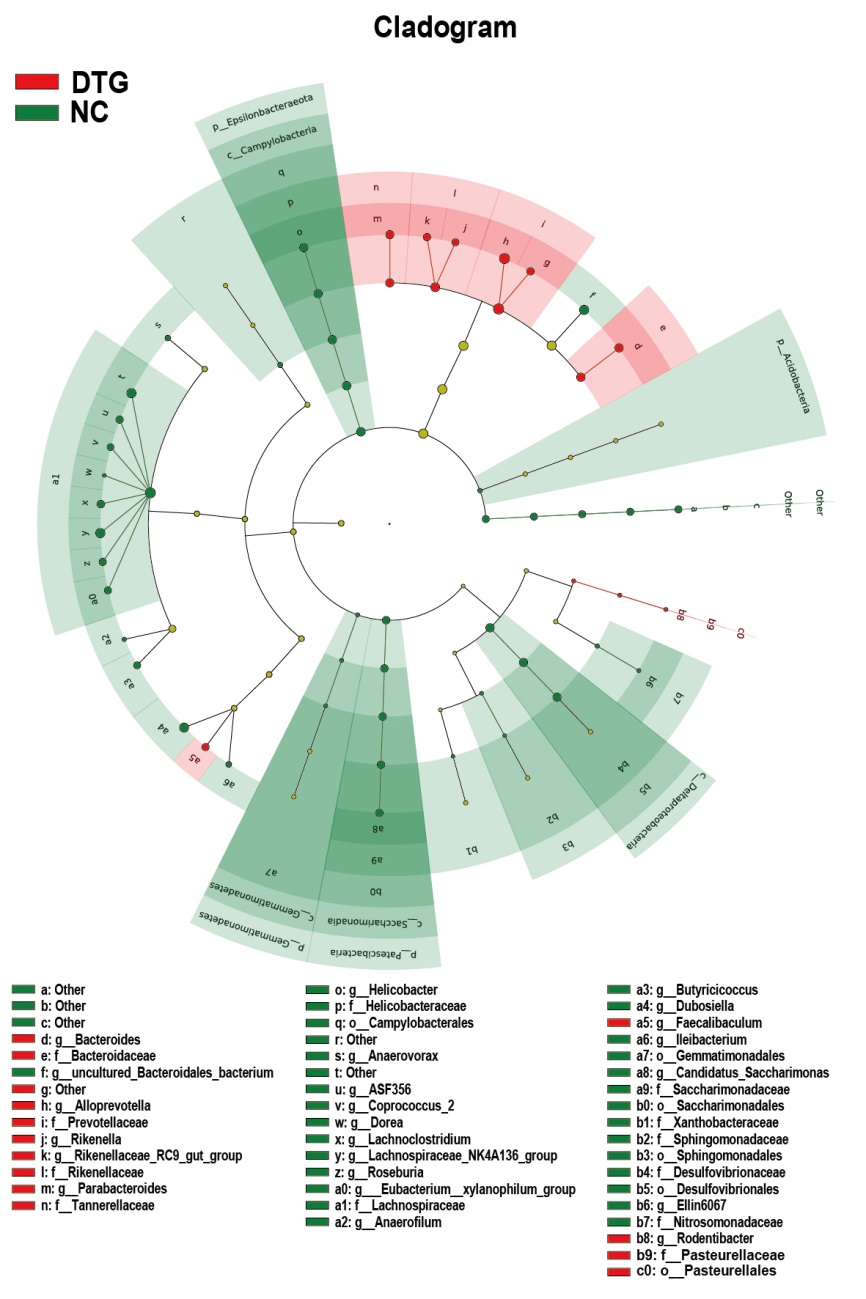

**Supplementary Figure 4. Cladogram of linear discriminant analysis effect size (LEfSe) showing the microbiota with different abundances between DTG and NC groups.**

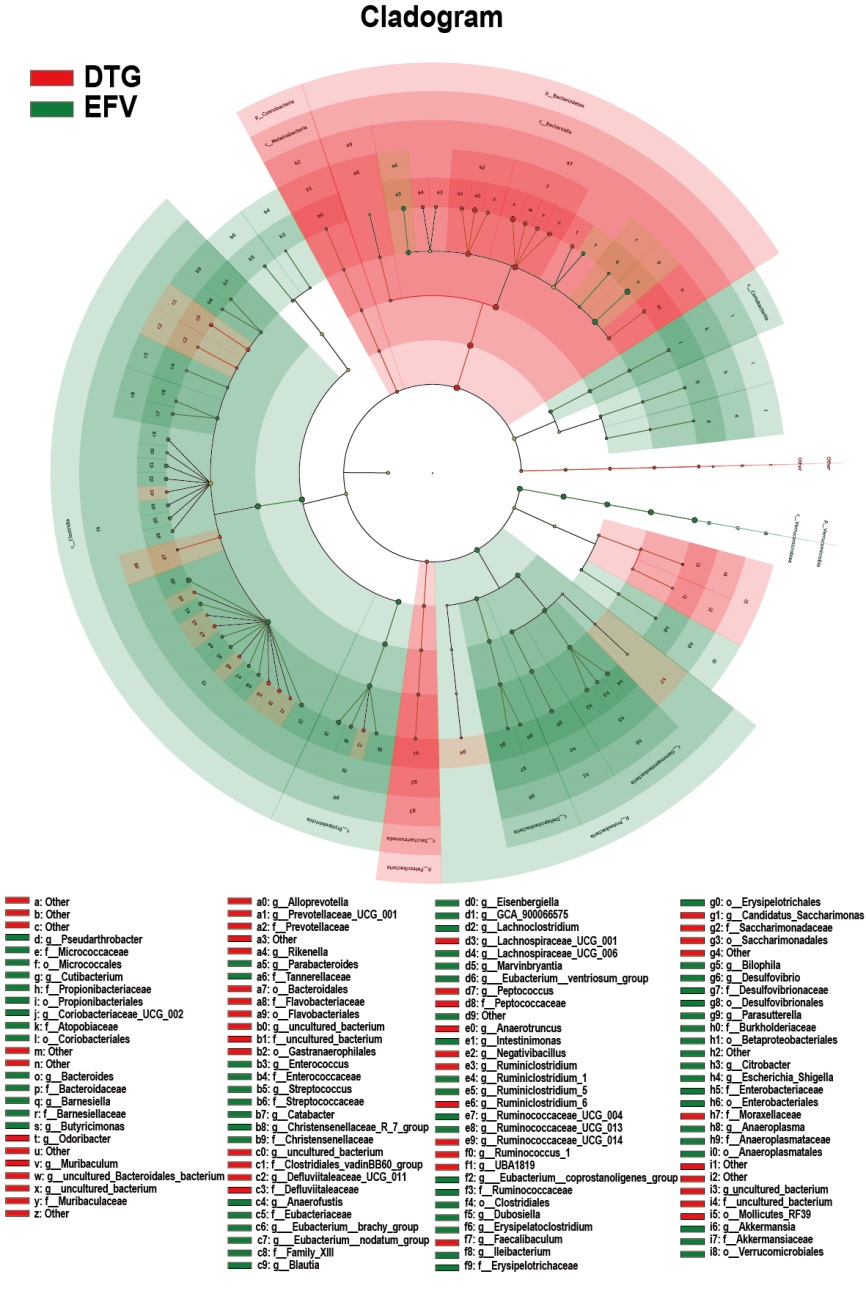

**Supplementary Figure 5. Cladogram of linear discriminant analysis effect size (LEfSe) showing the microbiota with different abundances between EFV and DTG groups.**
